# Epileptiform activity and seizure risk follow long-term non-linear attractor dynamics

**DOI:** 10.1101/2024.07.26.605279

**Authors:** Richard E Rosch, Brittany Scheid, Kathryn A Davis, Brian Litt, Arian Ashourvan

## Abstract

Many biological systems display circadian and slow multi-day rhythms, such as hormonal and cardiac cycles. In patients with epilepsy, these cycles also manifest as slow cyclical fluctuations in seizure propensity. However, such fluctuations in symptoms are consequences of the complex interactions between the underlying physiological, pathophysiological, and external causes. Therefore, identifying an accurate model of the underlying system that governs the multi-day rhythms allows for a more reliable seizure risk forecast and targeted interventions. To achieve this goal, we adopt the Hankel alternative view of Koopman (HAVOK) analysis to approximate a linear representation of nonlinear seizure propensity dynamics. The HAVOK framework leverages Koopman theory and delay-embedding to decompose chaotic dynamics into a linear system of leading delay-embedded coordinates driven by the low-energy coordinate (i.e., forcing). Our findings reveal the topology of attractors underlying multi-day seizure cycles, showing that seizures tend to occur in regions of the manifold with strongly nonlinear dynamics. Moreover, we demonstrate that the identified system driven by forcings with short periods up to a few days accurately predicts patients’ slower multi-day rhythms, which improves seizure risk forecasting.

## Introduction

Many biological processes exhibit endogenous cyclic behavior at different time scales, ranging from faster circadian rhythms to slower weekly or monthly cycles ^1;2^. Circadian rhythms, which repeat every 24 hours, are observed in several physiological processes, such as sleep cycles and gene expression patterns ^2^. Similarly, slower multi-day cycles are observed, including cardiac ^3^ and menstrual cycles ^4^. There has been long-standing observations of temporal patterns at similar time scales in pathological processes across various disorders, including immunological and cardiological, ^5;6^, and mental health disorders ^7;8^. Indeed, cyclic patterns were once considered so essential to certain paroxysmal disorders, that their terminology reflects this. For example, the term “lunatic” originates from the belief that epileptic seizures were caused by lunar cycles ^9^. Seizure cycles have long been described, and more recently validated quantitatively using seizure diary data ^10^ and long-term electroencephalographic recordings, e.g. from implantable neurostimulation devices such as Neuropace’s Responsive Neurostimulator (RNS) device ^11;12;13;14;15^. In addition to seizures, other markers of pathophysiological cortical excitability, such as interictal epileptiform activity (IAE) have also shown multiday cycles. There is a reported phase relationship between cycles estimated from both physiological and pathophysiological measures, suggesting that both may reflect shared underlying causes ^16;10^.

Cyclical physiological patterns often emerge from internal ‘clock’ like mechanisms, further shaped by external input. For example, diurnal variation in the hypothalamic-pituitary-adrenal axis is powerfully entrained by light exposure ^17^ but shows cyclical behavior even in the absence of external environmental cues. The relationship between internal drivers and external influences is less well understood for multi-day cyclical patterns. While lunar entrainment of seizure cycles was once a hypothesis, a closer examination of RNS’s interictal epileptiform activity (IEA) cycles and circular statistics has revealed no evidence supporting it ^15^.

Despite our limited mechanistic understanding, multi-day cycles have shown some promise in assessing and forecasting seizure risk for individual patients ^16;18^. However, accurately forecasting seizure cycles from prior data over the long term is particularly challenging due to the irregular changes in the cycles themselves over time. Such dynamic changes evolving over several temporal scales are not well captured by current modeling approaches. Therefore, developing a principled framework for generative modeling of these seemingly chaotic dynamics that go beyond statistical descriptions of limited observed data could improve our understanding of the underlying causes and enable more reliable forecasts of impending seizure events.

Modeling time-varying expression of pathophysiology using a dynamical systems framework has been used to represent e.g. dynamic patterns observed in psychopathology ^19^. Such a framework offers model-based predictions and simulations of control-theoretic interventions, such as drugs or neuromodulation therapy. However, the approach relies on prior knowledge about the system’s underlying dimensions and equations, which are typically not known for complex dynamical pathophysiology such as epilepsy.

Recent advances in data-driven system identification methods, such as the sparse identification of nonlinear dynamical systems (SINDy) ^20^, provide a reliable tool for discovering the equations and normal forms ^21^ of real-world dynamics. Such simplified representations of the high-dimensional nonlinear or chaotic dynamics are beneficial, as they provide valuable information about the system’s important macroscopic behavior, such as the emergence of bifurcations ^22^ or switching phenomena ^23^.

The occurrence of individual seizures is likely caused by multiple factors, reflecting the dynamics of the high-dimensional system of physiological, pathophysiological, and environmental causes. The temporal profile of seizure risk can be approximated with some accuracy from individual physiological measurements ^10;18^, but these do not reflect the complexity of the full underlying dynamics. However, we can reconstruct a diffeomorphic attractor from a single measurement to the underlying measured attractor based on Taken’s seminal embedding theorem ^24^. Taken’s theorem implies that the attractors’ full dynamics and important information regarding their topology can be uncovered by delay-embedding a single relevant measurement. More recently, a framework combining the delay-embedded coordinates and dynamic mode decomposition – Hankel Alternative View of Koopman (HAVOK) – was shown to allow the decomposition of nonlinear chaotic dynamics into a linear dynamical system with intermittent forcing captured by the low-energy delay coordinates ^23^. Increases in forcing were shown to mark periods with high nonlinearity followed by switching and bursting behavior *in silico* as well as in real-world examples. For instance, increased forcing predicts the switching between the lobes in the Lorenz attractor, changes in the earth’s magnetic field, and measles outbreaks ^23^.

In this study, we utilized the delay-embedding approach to investigate the topology of attractors underlying multi-day seizure cycles in patients with RNS implants. Our results indicate that a linear model driven by higher frequency forcing, using delay-embedded coordinates, can accurately predict the future trajectory of IEA-count time series as well as the occurrence of ‘long events’ (LEs). These are automatically detected seizure-like electrocorticography patterns, the detection of which is individually tuned by the clinicians to reflect patient-specific seizure patterns. Additionally, we observed that an increase in forcing drives the system into state-space regions marked by non-linear dynamics and increased seizure likelihood. Notably, our findings demonstrate that the forcing can accurately forecast impending seizures for up to several days in most patients, providing a data-driven approach that offers a mechanistic view of the chaotic dynamics of seizure cycles. The model output may represent an accurate model-based biomarker for impending seizures in patients undergoing long-term electrophysiological monitoring.

## Results

We hypothesized that the multi-day cycles in hourly IEA counts of patients undergoing long-term electrocorticography monitoring through an RNS device are the reflection of an underlying nonlinear system with chaotic dynamics. In order to uncover the topology of the underlying attractor, we first applied a low-pass filter aiming to dampen circadian oscillations, with a 3-day cut-off period to the hourly IEA-count time series to reduce the power of the higher frequency circadian cycles (Figure 1a-b) – see Materials and Methods for more details on the rationale for low-pass filtering. Next, we constructed a high-dimensional (*d* = 100) delay-embedded matrix (i.e., Hankel matrix) by stacking time-shifted versions of the filtered hourly IEA-count time series. The delay-embedded matrix’s singular value decomposition (SVD) allows us to reduce the dimensions by extracting the *r* leading time-delay-coordinates. Specifically, in the context of a delay-embedded matrix *A*, the SVD can be expressed as *A* = *USV*^′^, where *U, S*, and *V*^′^ represent the orthogonal matrices that capture the dominant directions of variability in the rows and columns of *A*, and *S* denotes the diagonal matrix of singular values. Since *U* and *V*^′^ are unitary, their columns form a set of orthonormal vectors, which can be regarded as basis vectors. The columns of *V* are called the right singular vectors of *A* and provide the time series associated with each SVD component(Figure 1c). Taken’s embedding theorem allows us to reconstruct a *k*−dimensional attractor, diffeomorphic to the original attractor with box-counting dimension *d*_*a*_ (*k* > *d*_*a*_) from a single measurement. Therefore, in theory, a single measurement can uncover key features of the full dynamics of a complex biological system, which may reflect the dynamic trajectory underpinning variable seizure risk. Figure 1d demonstrates that the first three SVD coordinates reconstruct an attractor topology out of a single IEA time series for a sample patient.

**Figure 1:**
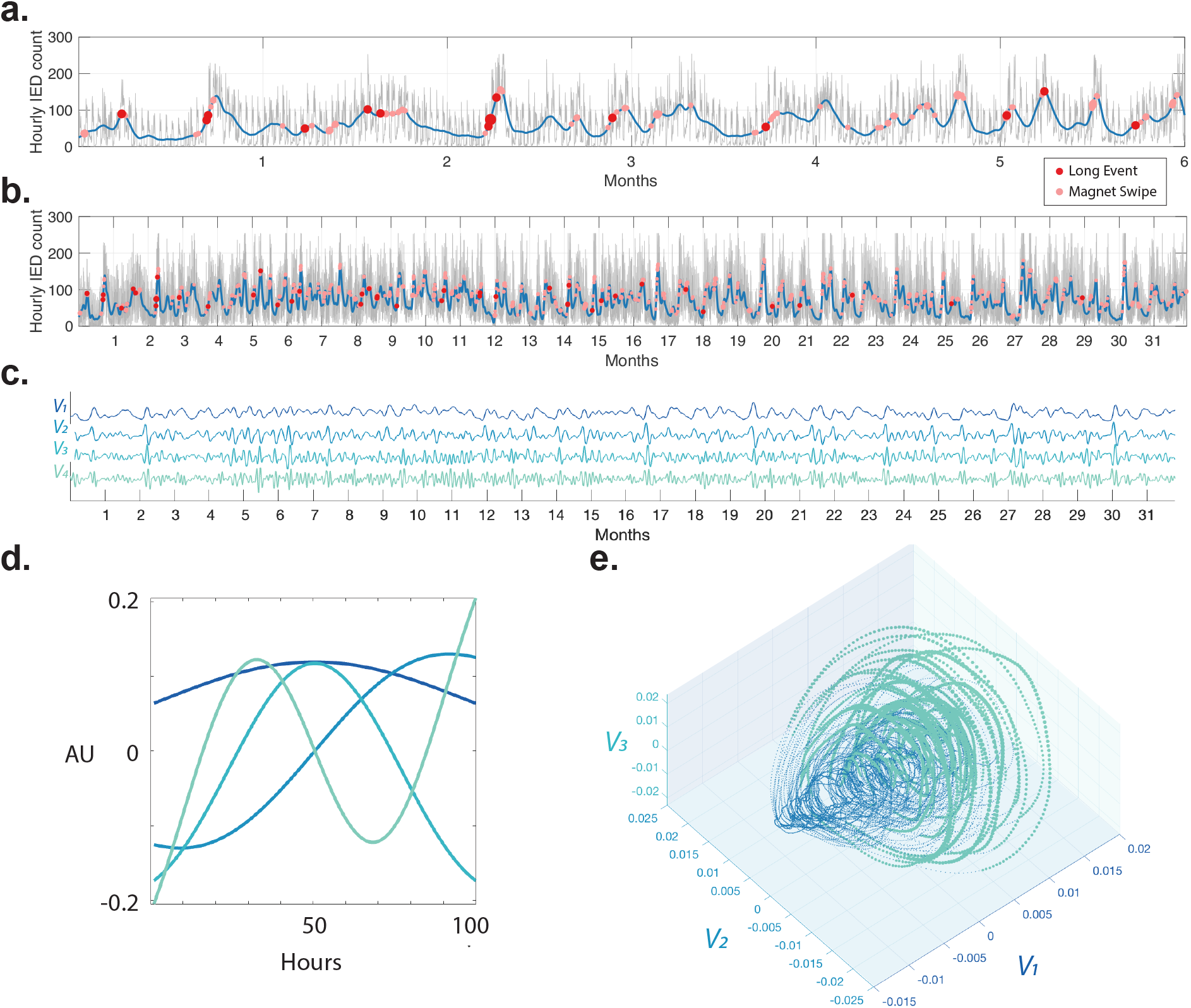
Delay-embedded coordinates of IEA-count time series reveal the topology of the attractor governing seizure risk. ***a***. IEA hourly count time series from a representative patient. The blue line indicates the low-pass (three-day) filtered time series. Red dots show clinical seizures labeled by the patient using magnet swipes. Pink dots show the LEs detected by the RNS device, and their size indicates their hourly count. ***b***. Same as ***a*** over a longer window (32 months). ***c***. The delay-embedded coordinate time series (*V* matrix columns) of the average IEA curve in panel ***b*** calculated using Singular Value Decomposition (SVD) of the delay-embedded matrix (i.e., Hankel matrix, see Materials and Methods for more detail). ***d***. The basis vectors (*U* matrix columns) corresponding to the delay-embedded coordinate time series in panel ***c. e***. Reconstructed attractor. Samples with larger amplitude (measured using line length) of the higher frequency delay-embedded coordinate (*V*_4_) oscillations are size- (amplitude) and color-coded (*V*_4_’s normalized line length > 0).

The dynamics of a chaotic attractor can be decomposed into a linear model in the *r* − 1 leading SVD coordinates forced by low-energy *r* coordinate (*V*_*r*_) ^23^. An increase in forcing precedes switching and bursting phenomena in several analytic and real-world systems. Accordingly, we hypothesize that seizures would reside in regions of the attractor with high non-linearity, preceded by increased forcing in the reconstructed system. We used the line-length measure (72-hour window) of *V*_*r*_ to quantify the gradual increase in the forcing’s amplitude envelope. Highlights in Figure 1d show the regions with high forcing (normalized *V*_*r*_’s line-length > 0). In this example, we selected *r* = 4 as the forcing for this three-dimensional system.

We use the device-labeled ‘long events’ (LEs) as a proxy measurement of ictal activity since these are tuned by the clinicians to identify patient-typical seizures and often represent electrographic seizures and correlate in frequency with seizure clinical frequency ^25^. The overlap between the patient-labeled seizures and the long event shows the close relationship between the two measures in Figure 1a. Figure 1e shows in the sample patient that the long events cluster over a part of the manifold preceded by increased forcing.

Next, we trained a classifier to detect LEs for every patient using six months of delay-embedded coordinates to test if the position in the phase space is predictive of seizure-like electrocorticography patterns (see materials and methods for details on the classification scheme). Our results show that in most patients, the higher frequency delay-embedded coordinates are predictive of LE occurrence (permutation test *n* = 100 *p* < 0.05, see Statistics and Reproducibility for details on the random null model, and the Materials and Methods section for details regarding the classification scheme) (Figure 2 and SI Figure 2.)

**Figure 2:**
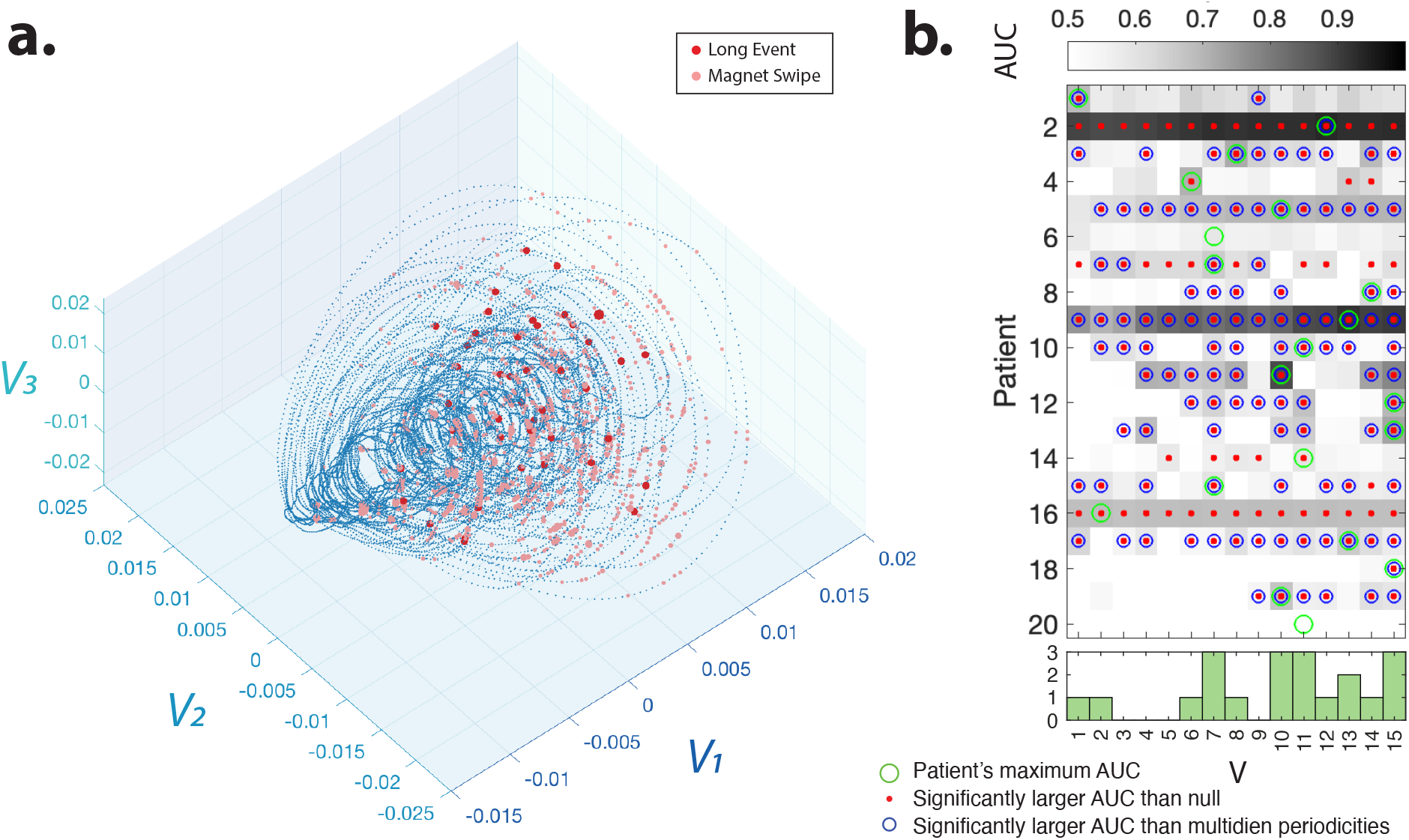
Delay-embedded IEA-count time series coordinates detect the seizure risk. ***a***. Patient-labeled seizures (red dots) and device-labeled automatically detected ‘long events’ (LEs, pink dots) overlap with the region of the manifold marked by the increased forcing in Figure 1e. ***b***. The mean daily area under the receiver operating characteristic curve (AUC) for detection of seizure risk (LEs) using different delay-embedding coordinates over 50 repetitions of the analysis (see Materials and Methods for classification details). The histogram (bottom panel) shows the number of patients with a peak AUC for a given SVD component. Coordinates derived from single SVD components predict LEs better than patient-specific multi-dien IEA cycles (see methods, peak paired t-test between the SVD component providing max. AUC (green circles), and two cycles; detection mean AUC (std) for SVD = 0.67 (0.13), for slow cyles = 0.56 (0.14), p=9.8e-04). Blue circles highlight each SVD that provides significantly better AUCs compared to multi-dien cycle features (two-sample *t*−test, *p* < 0.05, Bonferroni corrected for multiple comparisons). Red dots show the mean AUC values that are significantly higher than those of the random null detection (two-sample *t*−test, *p* < 0.05, Bonferroni corrected for multiple comparisons across patients and coordinates. See Statistics section for more details on the null and permutation test).

It has been demonstrated that seizures tend to occur during the ascending phase of multi-day cycles ^16^ and that the phase of multi-day cycles predicts seizure risk ^18^. Our results demonstrate that for most patients, delay-embedded coordinates provide significantly more accurate detection of long events (Figure 2a-b.) than phase and amplitude of the identified main two slow multi-day cycles (See SI Figure 1 and Materials and Methods for more details on identifying the multi-day cycles’ peaks). Interestingly, *V*_1_, which corresponds to the coordinate with the slowest oscillations, only predicts the LEs of a few patients. A subset of patients reliably self-reported seizures triggered externally by a magnetic wand. For these patients, delay-embedded coordinates also allow the detection of self-reported seizures, although different coordinates provide optimal detection across patients (SI Figure 3).

**Figure 3:**
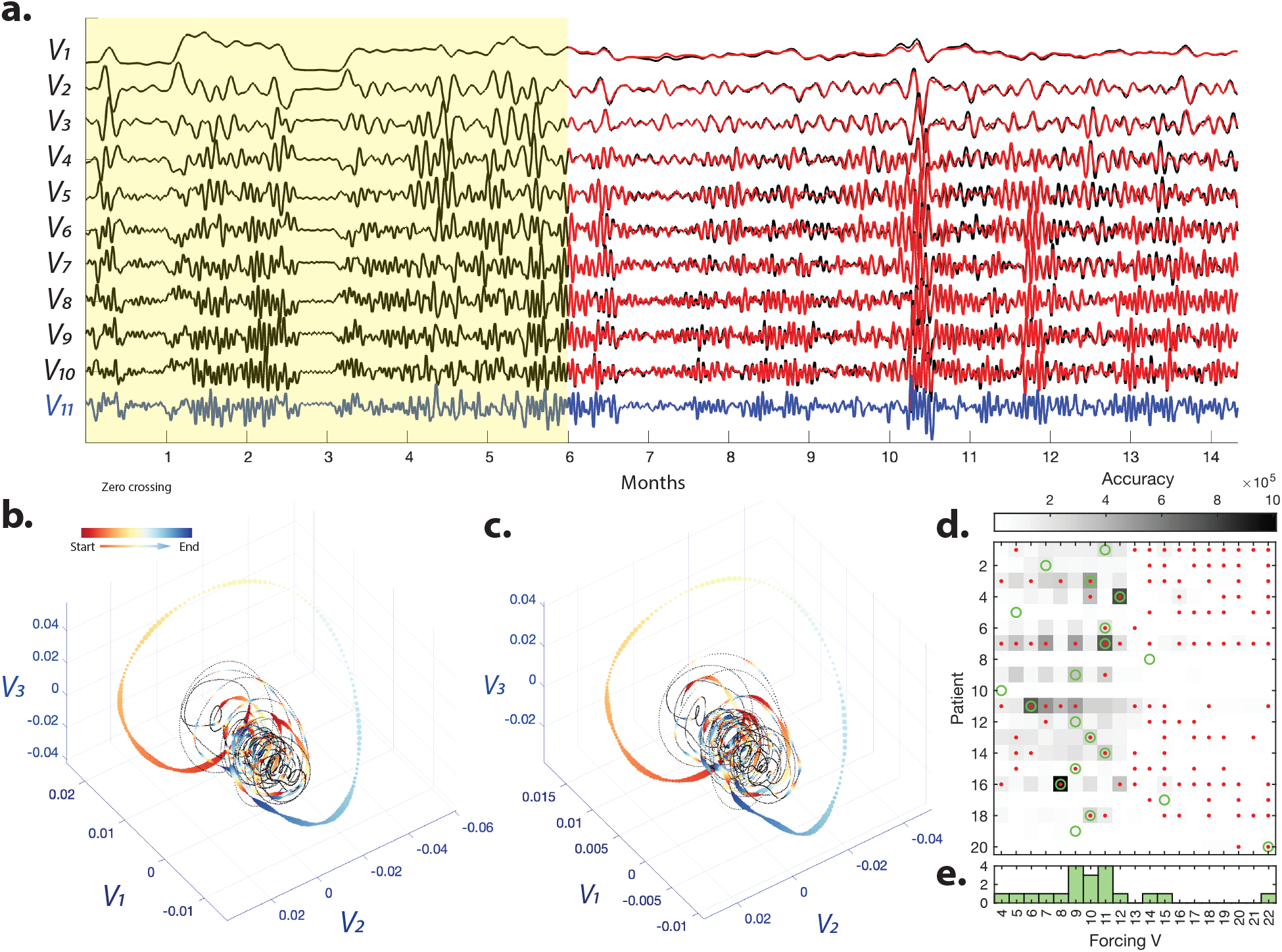
A linear model of the delay-embedded coordinates driven by higher frequency forcing (i.e., *V*_*r*_) accurately predicts the future trajectory of multidien IEA-periodicities. ***a***. Black line shows the *V*_1_ to *V*_11_ of a sample patient. We trained a *r* − 1 = 10 dimensional linear model using *V*_1_ to *V*_10_ and *V*_11_ (Blue line) as the forcing over a 6-month window (see Materials and Methods for per-processing details). The red lines show the predicted time series generated from the trained linear dynamical system driven by the known forcing (*V*_11_) over the test window (i.e. after the 6-month training window). The reconstructed embedded ***b***. and predicted attractors ***c***. from the test period. Instances of zero-crossing in the normalized line length of the forcing are color-coded. ***d***. Forecasting based on single SVDs performed better than forecasting based on multi-dien cycle features (peak AUC for single SVD forecasts versus multiday cycle features differ in 1-day forecast: for single SVD, mean peak AUC (std) = 0.65 (0.08), for multi-dien periodicities AUC (std) = 0.55 (0.1), paired t-test *p* =8.3698e-04; in 7-day forecast: single SVD mean peak AUC (std) = 0.61 (0.07), for multi-dien periodicities mean AUC (std) AUC = 0.55 (0.08, paired), paired t-test *p* =1.3280e-04). The plot shows the accuracy (the inverse of mean squared error) of the predicted leading coordinate (*V*_1_) time series for systems of different sizes with *V*_4_ to *V*_22_ as forcing. Red dots indicate significantly higher accuracy than those of the future trajectory predictions of the phase-randomized IEA-count null time series (*p* < 0.05, two-tailed, *n* = 50 iterations, FDR corrected for multiple comparisons across patients and delay-embedded coordinates). Green circles highlight the maximum accuracy for each subject. The histogram (bottom panel) indicates the number of subjects with maximum accuracy for each forcing.

Within the HAVOK framework, the low-energy forcing drives the leading delay coordinates. This implies that the forcing can be used to causally drive the dynamics and enable forecasting the trajectory of the full system. To test this hypothesis, we apply the HAVOK framework to train a linear model based on a six-month window. Next, we simulate the fitted linear system dynamics using the known future forcing time series (i.e., *V*_*r*_). As seen in Figure 3 in a sample patient, the simulated *V*_1_ to *V*_10_ time series using the *V*_11_ as the forcing closely match the delay embedded coordinates and reconstructs the embedded attractor. As seen in SI Figure 4, this forcing (*V*_11_) displays fat-tailed distributions corresponding to intermittent and rare events.

**Figure 4:**
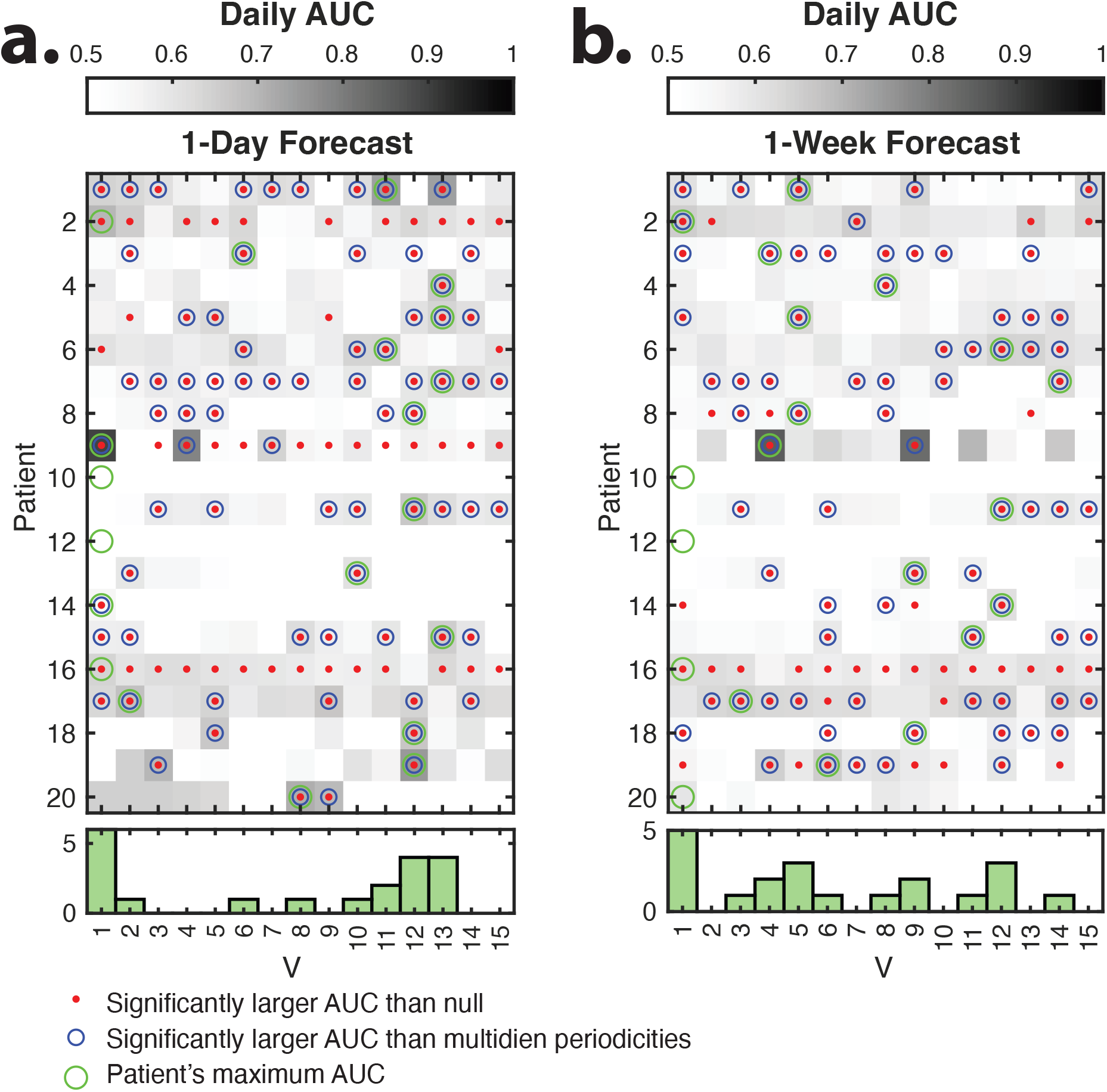
Delay-embedded coordinates of the IEA-count time series enable forecast of the seizure risk. The mean daily Area Under the Receiver Operating Characteristic Curve (AUC) for one day ***(a)*** and one week ***(b)*** forecast of seizure risk (long events) using different delay-embedding coordinates over 50 repetitions of the analysis (see Materials and Methods for classification details). Red dots show the mean AUC values that are significantly higher than those of the random null forecast (two-sample *t*−test, *p* < 0.05, Bonferroni corrected for multiple comparisons across patients and coordinates. See Statistics section for more details on the null and permutation test). Blue circles show the mean AUC values that are significantly (two-sample *t*−test, *p* < 0.05, Bonferroni corrected for multiple comparisons) higher than the AUC values calculated from the two slow peak features (i.e., their amplitude and phase). Green circles show the maximum mean AUC across all coordinates for each patient. Histograms (bottom panels) indicates the number of subjects with maximum AUC for each forcing

This close fit suggests that an intermittently forced linear model can accurately capture the chaotic dynamics underpinning ictogenicity and highlight the identified forcing’s potential for seizure risk forecast. Moreover, we quantified the accuracy – measured as the inverse of the mean squared error – of predicted trajectories of the leading coordinate (*V*_1_) in Figure 3d and the three leading delay-embedded coordinates of the IEA-count time series for all patients in SI Figure 5. As seen in Figure 3d, the accuracy of the predicted leading coordinate (*V*_1_) increases for higher dimensional linear systems, followed by a sharp drop in accuracy at the 12-dimensional system (*V*_13_ as forcing) for all patients. SI Figure 6 demonstrates that *U*_10_ to *U*_12_ have prominent 25-hour cycle peaks for almost all patients and *U*_13_ corresponds to faster cycles (period < 25 hours).

The delay-embedded coordinates and the forcing time series are extracted from the entire IEA-count time series using the singular value decomposition of the delay-embedded Hanekl matrix. Alternatively, we can extract the forcing time series of the test time series directly using the basis vector of the *U* matrix estimated from the six-month window. Specifically, it is possible to measure forcing (*V*_*r*_) from a streaming time series by convolution with the *r*_*th*_ mode (column *r* of matrix *U*). Our results show that similarly, the convolution-based estimated forcing accurately predicts trajectories of the three leading delay-embedded coordinates of the IEA-count time series (see SI Figure 7 for details). Like the above-mentioned results, the forcing corresponding to the daily cycles provides the most accurate predictions. Together, this close fit suggests that an intermittently forced linear regression model can accurately capture the chaotic dynamics underpinning ictogenicity and highlights the estimated forcing’s potential for seizure risk forecast.

Finally, to test the hypothesis that the delay-embedded coordinates enable the forecast of future seizure risk, we train a classifier using the time-shifted long event time series (1-to 7-day shifts) and measure the forecast accuracy. In most patients, delay-embedded coordinates forecast LEs daily risk in the next day to week significantly better than randomized nulls (*t*−test, *p* < 0.05, Bonferroni corrected for multiple comparisons across patients, 4). For the majority of patients, the highest AUCs for one-day forecast were achieved for faster delay-embedded coordinates (*V*_*r*_), *r* > 6 period length <= 40*h* 4a). Overall, this method performed better than forecasting based on the main two identified multi-day 4). Similar results are observed for longer 2-to 7-day forecasts of long events (SI Figure 8) and patient-labeled seizures (SI Figure 9). Together, these results validate the utility of the identified delay coordinates and the higher frequency forcing for improving the accuracy of the seizure risk forecast.

## Discussion

In this study, we have introduced a novel data-driven model-based approach to approximate the dynamics of a putative dynamical system driving slow multi-day rhythms in seizure risk in individual patients undergoing long-term electrocorticography monitoring for seizures. We aimed to reconstruct a linear dynamical system intermittently influenced by higher-frequency low-energy drivers based on the complex long-term IEA-count time series obtained from the RNS device. Our findings demonstrate the effectiveness of this framework in capturing the causal relationship between the higher frequency forcings and the slow multi-day oscillations and accurately predicting the system’s future trajectory based on the modeled dynamics. By leveraging the high-frequency delay-embedded coordinates as drivers, we successfully drive the slow multi-day oscillations *in silico* and accurately reconstruct the future trajectory of the full system in most patients, shedding light on the complex interactions underlying seizure propensity dynamics.

Importantly, our results bear clinical significance as they showcase the potential of our approach to enhance the detection and forecasting of impending seizures over extended periods with patient-specific data-driven models. By leveraging the insights gained from our model, we demonstrate improved capabilities in identifying and anticipating seizure events over several days, providing valuable information for clinicians and patients. This work establishes a framework for quantitatively characterizing relationships between circadian and slow multi-day rhythms, seizure risk dynamics, and external drivers. This holds promise for developing more reliable seizure risk forecasting models and personalized interventions based on patient-specific physiological recordings.

Our work highlights the advantages of adopting a linear model to capture the interactions between slower modes driven by faster modes and the low-energy external forcing compared to the conventional wavelet-based analysis of the IEA-count time series. This novel approach offers several key benefits. Firstly, it provides a data-driven and model-based improvement on established methods, currently based on the subjective selection of peaks in the IEA wavelet power spectra (e.g., ^16;26^), which often relies on arbitrary criteria to identify the slow oscillatory modes. Secondly, unlike our model-based framework, wavelet analysis fails to unveil the intricate relationship between the slower and faster dynamics. In contrast, our proposed linear model effectively uncovers this crucial connection, shedding light on the underlying mechanisms driving the observed phenomena. SI Figure 1 illustrates how spectral analysis of the IEA-count time series wavelet decomposition can provide additional insight into the frequency of amplitude modulations at various scales. However, these wavelet analysis results remain purely descriptive, lacking our linear model’s explanatory power and predictive capabilities.

Previous studies have highlighted the clinical significance of IEA cycles, specifically the correlation between the phase of multidien periodicities and seizure risk ^16;14^. However, these proof-of-concept studies primarily rely on non-causal filtering of the IEA-count time series (e.g., ^18^), limiting their practical application for real-time seizure risk forecasting. The critical advantage of our findings and framework lies in its ability to directly predict the emergence of slow oscillations from their higher-frequency drivers, such as daily cycles, using the modeled system. This approach eliminates the lengthy delays introduced by causal low-pass filtering when forecasting the impending increase in IEA counts. Instead, our method leverages faster oscillations, which theoretically exhibit shorter delays, resulting in more timely and accurate seizure risk predictions.

Posing the multi-scale IEA cycles as outputs of intermittently forced attractors presents a novel avenue for comprehending the drivers of ictogenicity. By examining the estimated slow coordinates and forcing, we can uncover connections to physiological factors (e.g., metabolic or hormonal rhythms ^27^) or pathophysiological processes. Furthermore, our model-based approach provides a framework for testing hypotheses regarding external or internal factors. For instance, we can model and predict the impact of external interventions, such as drugs or neurostimulation treatments, on the impulse response of the modeled system and IEA cycles. Similarly, future research could aim to enhance prediction accuracy by explicitly adding known external stressors, such as weekly stressors, to the model.

Interestingly, our findings indicate that changes in the circadian IEA cycles and the high-dimensional model of the IEA-count time series offer the most accurate forecast of future IEA-count trajectories for most patients. However, most patients’ circadian IEA cycles were not the most predictive of future seizure risk. Instead, other slower oscillations demonstrated better accuracy in forecasting seizure risk in several patients. These results suggest that while circadian forcing can effectively predict the future of slow oscillations, the clinical manifestation of seizures may be better explained and mechanistically linked to a few or multiple slower coordinates within the system.

One limitation of our study is that we do not consider the impact of electrical stimulations delivered by the responsive neurostimulation devices on the IEA-count cycles. Recent studies have shown that neuromodulation can alter cycles ^28^, and stimulation can suppress EEG signal features ^29;30^. Nevertheless, our framework allows accounting for this by modeling the responsive neurostimulation and its effects on the slow cycles as a closed-loop system with feedback stimulation. There are other important caveats to consider regarding our findings, such as the unaccounted changes in the detection and stimulation parameters of the RNS device, as well as the use of long events and patient-labeled seizures as a proxy for clinical seizures, which have previously been shown to correlate with clinical seizure frequency ^25^. Additionally the small cohort sample size may limit generalizability. Nonetheless, our study contributes to the expanding knowledge base on slow multi-day rhythms in epilepsy, providing evidence for the clinical significance of accurately modeling and predicting seizure dynamics. The implications of our findings extend beyond theoretical implications, holding promise for advancements in seizure management and improving patient care.

## Materials and Methods

### Patient population information

We conducted a retrospective analysis of longitudinal hourly counts of detected interictal epileptiform activity (IEA) based on long-term electrocorticography recordings obtained from a subset of 20 out of 28 patients with drug-resistant epilepsy who underwent implantation of the RNS System (NeuroPace, Inc., Mountain View, CA) at the Hospital at the University of Pennsylvania. Our inclusion criteria required patients to have a minimum of one year of IEA recordings, resulting in the exclusion of eight patients. The 20 patients included in this study were implanted with the RNS device between August 2015 and January 2022. Before participating in this study, all patients provided written informed consent following the Institutional Review Board of the University of Pennsylvania.

### RNS IEA-count data preprocessing

In addition to providing real-time responsive neurostimulation, the RNS system keeps track of the hourly number of interictal epileptiform activity (IEA). To ensure the early and effective detection of seizures, the clinicians manually tune the detection parameters of the RNS device or patients’ office visits. Consequently, the detected hourly IEA counts can vary notably pre and post-office visits. To address this issue, we normalized (z-score) the IEA-count time series between each office visit. We also remove the first five months of data to account for implantation effects and other factors that contribute to high variability commonly observed early in the IEA-count time series.

### Identifying multi-day cycles’ frequency peaks

We employed a methodology similar to that described in Baud et al. (2018) to extract multi-day cycles from the IEA hourly counts ^16^. In brief, for each patient, we applied a wavelet transform to the IEA-count time series and identified the peaks in the average power across time scales (peaks in the periodogram) ranging from 3-45 days. Specifically, we selected two multi-day peaks with the largest powers, one between 3-11 days and the slowest between 7-45 days.

### Hankel alternative view of Koopman (HAVOK) analysis

We followed the Hankel alternative view of Koopman (HAVOK) methodology and analysis developed by Brunton et al. ^23^. Specifically, we create a delay-embedding (Hankel) matrix from the IEA-count time series by stacking 100 delayed (each row one time point delayed compared to the previous row). We tuned the stacking parameter to capture the dominant frequencies in the time series. Namely, too small or too big stacking values can lead to loss of sensitivity to low or high-frequency modes, respectively. Next, to find the eigen-delay-embedding coordinates, we applied Singular Value Decomposition (SVD) to the Hankel matrix. The Hankel matrix, denoted as *A*, can be decomposed using Singular Value Decomposition (SVD) into the form *A* = *USV*^′^, where *U, S*, and *V*^′^ represent orthogonal matrices that capture the primary directions of variability in the rows and columns of *A*. The matrix *S* contains the singular values along its diagonal. Importantly, the columns of the unitary matrices *U* and *V*^′^ form sets of orthonormal vectors, which can be interpreted as basis vectors. Specifically, the columns of matrix *V* correspond to the right singular vectors of *A*, providing the time series associated with each SVD component.

SI Figure 10 shows the first five columns of *U*. Note that although, as expected, *U*_1_ to *U*_5_ captures the lowest to faster oscillations, circadian cycles are (24-hour cycles) present in all the basis vectors. This is due to the relatively high power of circadian cycles in the IEA-count time series. As mentioned, the HAVOK framework aims to decompose the chaotic and non-linear time series into a linear system forced by low-energy, high-frequency modes. However, the strong presence of high-power circadian IEA-count cycles poses an issue for this framework and can lead to the aforementioned mixing of frequencies in the basis vectors. Moreover, it has been observed that models are more accurate and predictive when the basis vector *U* resembles polynomials of increasing order, as shown in Figure 1d.

To address this issue, we low-pass (3 days) filtered the IEA-count time series using a non-causal filter (FIR filter, Order=100). Figure 1d. shows a sample patients’ *U*_1_ to *U*_4_ of the low-pass filtered time series, which shows these basis vectors now only represent the lowest frequencies. The main frequency peak of the *U* basis vectors identified using Fast Fourier transformation also reveals the monotonous decrease in the main frequency of *U*_*r*_ basis vectors for higher values of *r* (SI Figure 6). These observations were the basis for choosing the low-pass filtering reprocessing step.

Next we construct a linear model of the first *r* − 1 variables of *V* driven by *V*_*r*_:

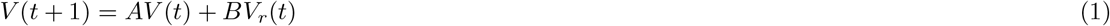

Here *V* (*t*) = [*V*_1_, *V*_2_, …, *V*_*r*−1_]^*T*^ is the *r* − 1 leading delay-embedded coordinates time series at time point *t, V*_*r*_(*t*) act as the input or forcing, and *A* and *B* are the system and input matrices, respectively. We estimated the system and input parameters from a six-month IEA-count time series window. Specifically, we used scripts provided by ^20^, which fit a discrete linear system by solving a least-squares problem. Next, we simulate the full future trajectory of the modeled system (i.e., *V* (*t*)) using the *V*_*r*_(*t*) time series after the six-month point. We measured the accuracy of the predicted V(t) separately for each dimension using the inverse of the mean squared error. We reported the accuracy of the predicted first leading three dimensions, *V*_1_,*V*_2_, and *V*_3_, in SI Figure 5. We also explored the convolution of the basis vector *U*_*r*_ estimated from the first six months with the full IEA-count time series to extract the forcing (*V*_*r*_) after the six-month time points. SI Figure 7 shows the accuracy of the predicted leading system dimensions using the convolution-based estimated forcing.

### Seizure risk detection and forecast

We examined two identified slow multi-day peaks to investigate the utility of amplitude and phase features in detecting and forecasting seizure risk. For the detection analysis, we applied a non-causal filter (Finite Impulse Response (FIR) filter, order=100) to the preprocessed IEA-count time series centered at the slow multi-day peak frequency *±* one day. However, for the forecast analysis, we leveraged a causal filter and shifted the long IEA events (or patient-labeled seizures) by *x* days (ranging from *x* = 1 to 7) to assess the predictive power of features in real-time scenarios. We then calculated the instantaneous phase of the filtered time series using the Hilbert transform.

Additionally, we explored the performance of delay-embedded coordinates and forcing in seizure detection and forecasting. In the detection analysis, we used the delay-embedded coordinates time series *V*_1_, …, *V*_*r*_ and their instantaneous phase as features to predict seizure risk. In the forecast analysis, apart from shifting the seizure labels, we directly extracted the forcing time series *V*_*r*_ from the IEA-count time series using convolution with *U*_*r*_. We exclusively utilized the convolution-derived *V*_*r*_ time series and its instantaneous phase as features.

We used slow multi-day and delay-embedded coordinates features separately to train a regression model using a bagging ensemble of decision trees (minimum leaf size = 8) and a support vector machine (SVM) regressor. After removing an initial 5-month data window (to account for early implant effects), we trained the model using the first six months of patients’ recording. We tested the model’s accuracy using the Area under the Receiver operating characteristic Curve (AUC) on the remaining time series. In addition to hourly results, we also examined the daily accuracy by training the model using the average feature values and the total sum of long IEA event counts (or patient-labeled seizures) over 24 periods.

### Statistics

We tested the statistical significance of the detection and forecast AUC values against the null model results. Specifically, we randomly shuffled the long IEA-count time series to train and test the accuracy of the abovementioned model. We created distributions of empirical and null AUC values by repeating the analysis 50 times. We used *t*−test (*p* < 0.05, two-tailed, FDR corrected for multiple comparisons across patients and delay-embedded coordinates) to assess whether the AUC values are significantly larger than the null model’s. Similarly, we compared the performance of the slow multi-day features against the delay-embedded features by creating two distributions of AUC values (*n* = 50 iterations) and the subsequent *t*−test (*p* < 0.05, two-tailed, FDR corrected for multiple comparisons across patients and delay-embedded coordinates).

We employed statistical testing to determine the significance of the accuracy in projected trajectories of the modeled linear system, as described in the “Hankel alternative view of Koopman (HAVOK) analysis” section of the Materials and Methods. This testing allowed us to establish whether the predicted trajectories performed significantly better than what would be expected by chance alone. Specifically, we repeatedly (*n* = 50 iterations) trained and tested the linear model on the phase-randomized IEA-count time series to create a distribution of null accuracy results. As mentioned above, we calculated the prediction accuracy of leading delay-embedded coordinates *V*_1_, *V*_2_, and *V*_3_ separately using the inverse mean squared error. Finally, we used *t*−tests (*p* < 0.05, two-tailed, FDR corrected for multiple comparisons across patients and delay-embedded coordinates) to identify significantly high accuracy projections.

## Data Availability

Full patient datasets are subject to patient confidentiality. Summary statistics and limited anonymized patient metadata can be made available upon reasonable request.

## Code Availability

The custom scripts are available at https://github.com/b-schd/RNS_HAVOK/.

## Acknowledgements

R.E.R. received funding from the Wellcome Trust (209164/Z/17/Z) and from the European Union’s Horizon 2020 research and innovation program under Specific Grant Agreement No. 945539 (Human Brain Project SGA3). A.A. was supported by the startup funding from the Department of Psychology at the University of Kansas).

## Author Contributions

R.E.R.: Conceptualization of this study, Methodology, Writing. B.S.: Conceptualization of this study, Data Preprocessing, Methodology, Data Analysis, Writing. B.L.: Conceptualization of this study, Writing. A.A.: Conceptualization of this study, Methodology, Data Analysis, Writing.

## Competing Interests

The authors declare no competing interests.

## Supplementary Figures

**Figure 1:**
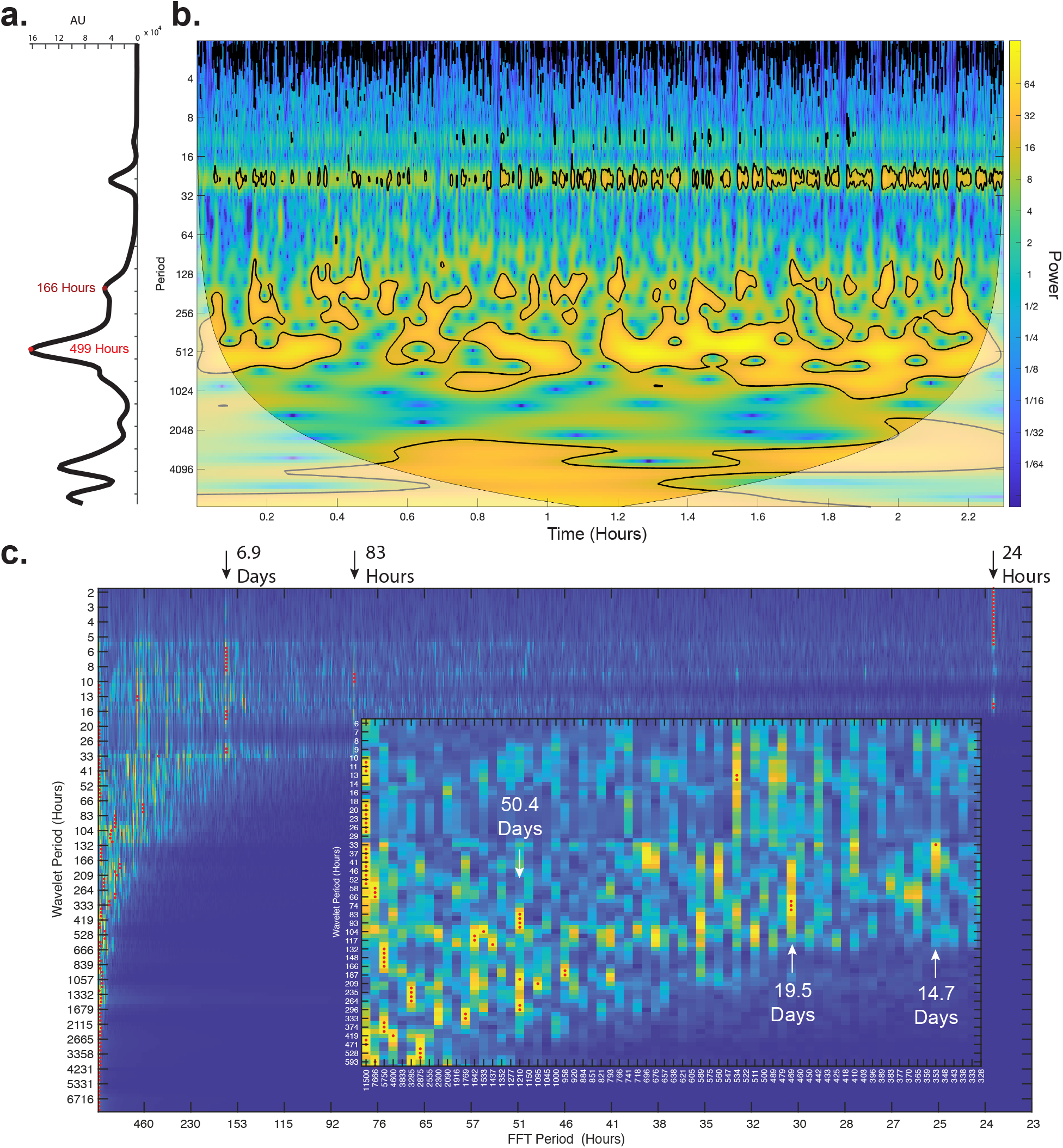
Identifying the slow multi-day peaks. The periodogram ***a***. and wavelet ***b***. decomposition of a sample patient’s IEA-count time series. The periodogram in panel ***a*** is created by summing the power of the wavelet decomposition over all time points for each scale (i.e., period). The two identified multi-day peaks are indicated on the plot. ***c***. Fast Fourier Transform (FFT) decomposition of wavelet time series for each scale. The red dots show the identified peak frequency. We highlighted the period of several dominant power amplitude-modulating frequencies with arrows.

**Figure 2:**
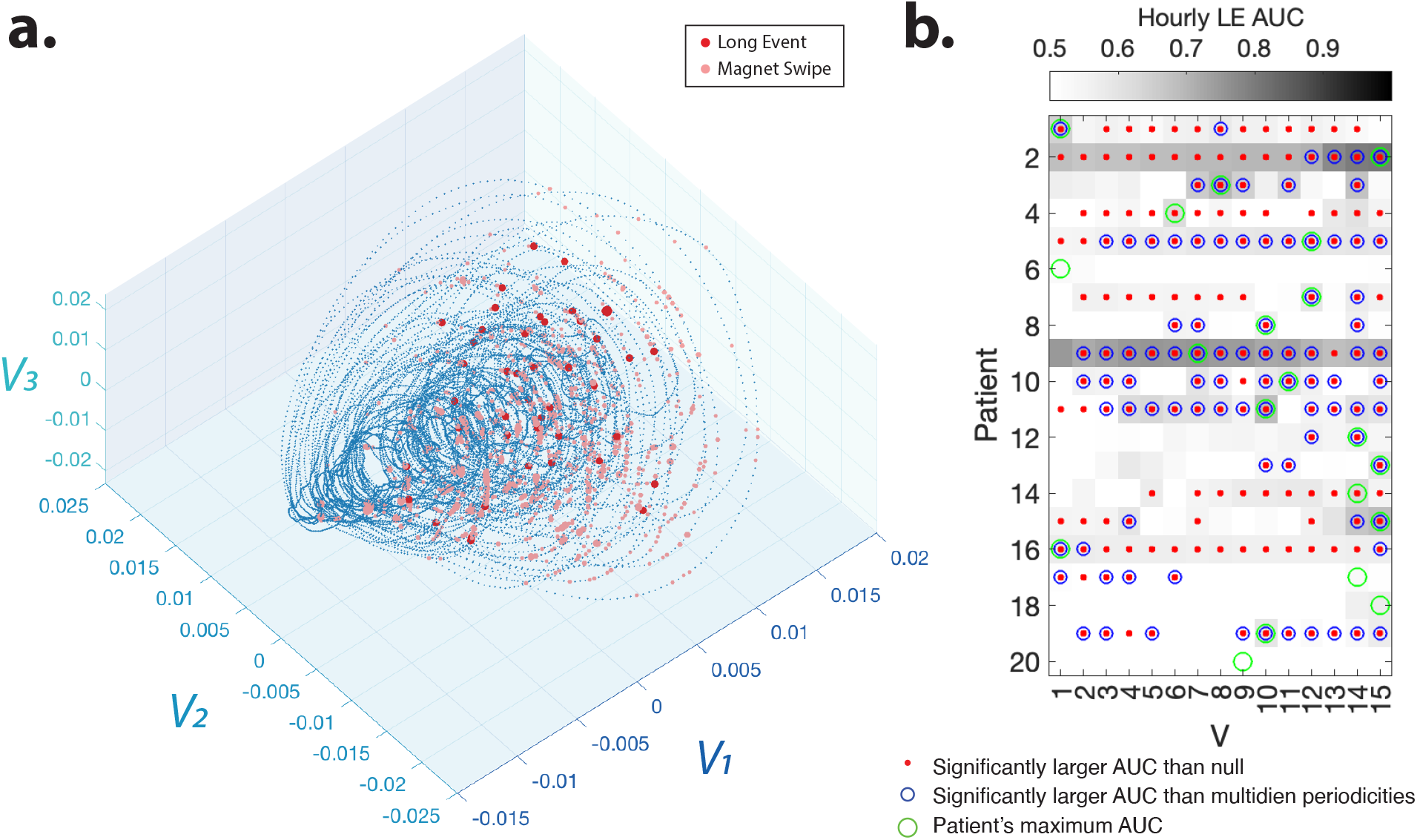
Delay-embedded IEA-count time series coordinates detect the seizure risk. ***a***. Patient-labeled seizures (red dots) and device-labeled long IEA events (pink dots) overlap with the region of the manifold marked by the increased forcing in Figure 1e. ***b***. The mean hourly area under the receiver operating characteristic curve (AUC) for detection of seizure risk (long IEA events) using different delay-embedding coordinates over 50 repetitions of the analysis (see Materials and Methods for classification details). Red dots show the mean AUC values that are significantly higher than those of the random null detection (two-sample *t*−test, *p* < 0.05, Bonferroni corrected for multiple comparisons across patients and coordinates. See Statistics section for more details on the null and permutation test). Blue circles show the mean AUC values that are significantly (two-sample *t*−test, *p* < 0.05, Bonferroni corrected for multiple comparisons) higher than the AUC values calculated from the two slow peak features (i.e., their amplitude and phase). Green circles show the maximum mean AUC across all coordinates for each patient.

**Figure 3:**
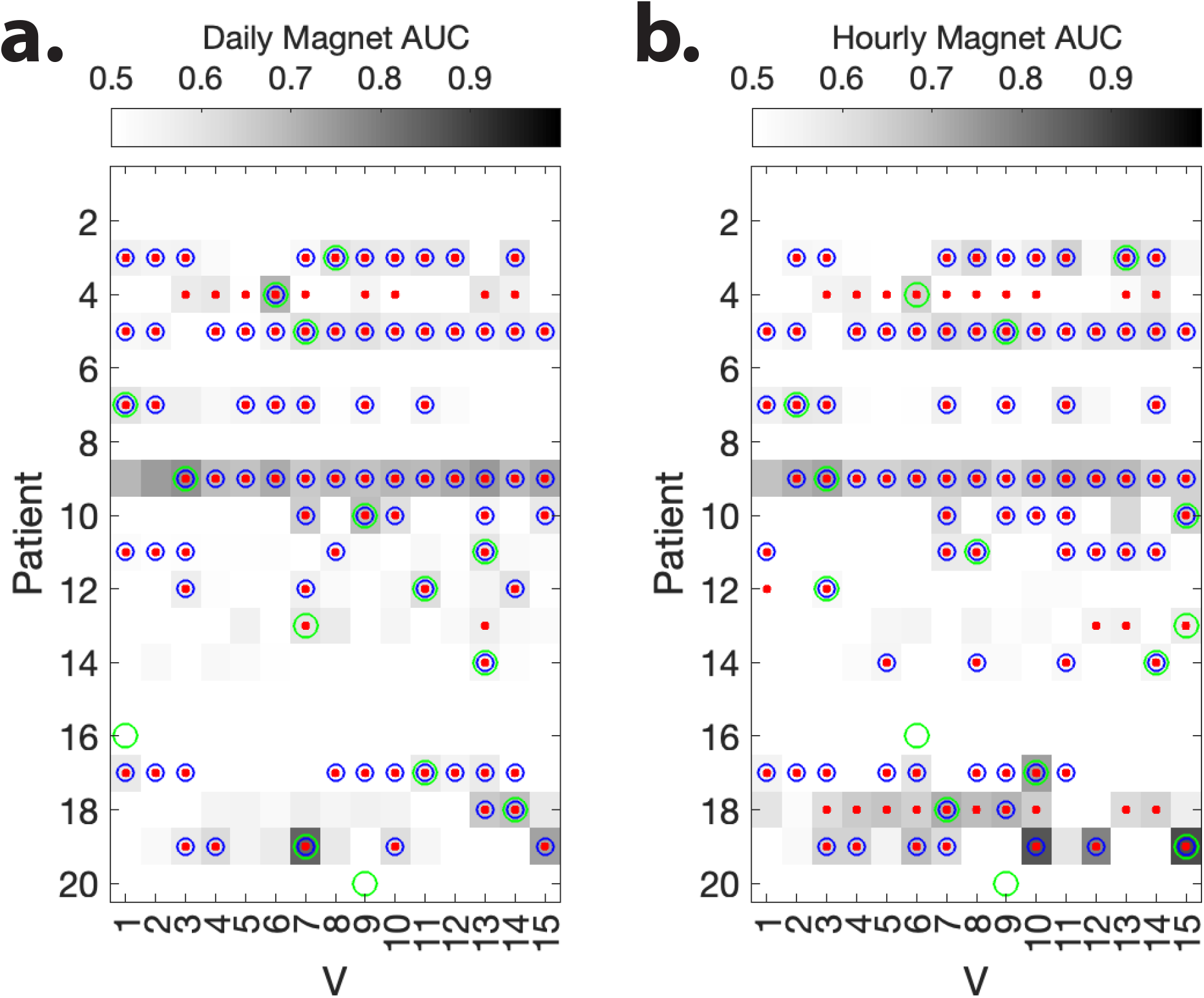
Delay-embedded coordinates of the IEA-count time series detect the patient-labeled seizure risk. The mean daily ***(a)*** and hourly ***(b)*** Area Under the Receiver Operating Characteristic Curve (AUC) for detection of patient-labeled seizure risk using different delay-embedding coordinates over 50 repetitions of the analysis (see Materials and Methods for classification details). Red dots show the mean AUC values that are significantly higher than those of the random null detection (two-sample *t*−test, *p* < 0.05, Bonferroni corrected for multiple comparisons across patients and coordinates. See Statistics section for more details on the null and permutation test). Blue circles show the mean AUC values that are significantly (two-sample *t*−test, *p* < 0.05, Bonferroni corrected for multiple comparisons) higher than the AUC values calculated from the two slow peak features (i.e., their amplitude and phase). Green circles show the maximum mean AUC across all coordinates for each patient.

**Figure 4:**
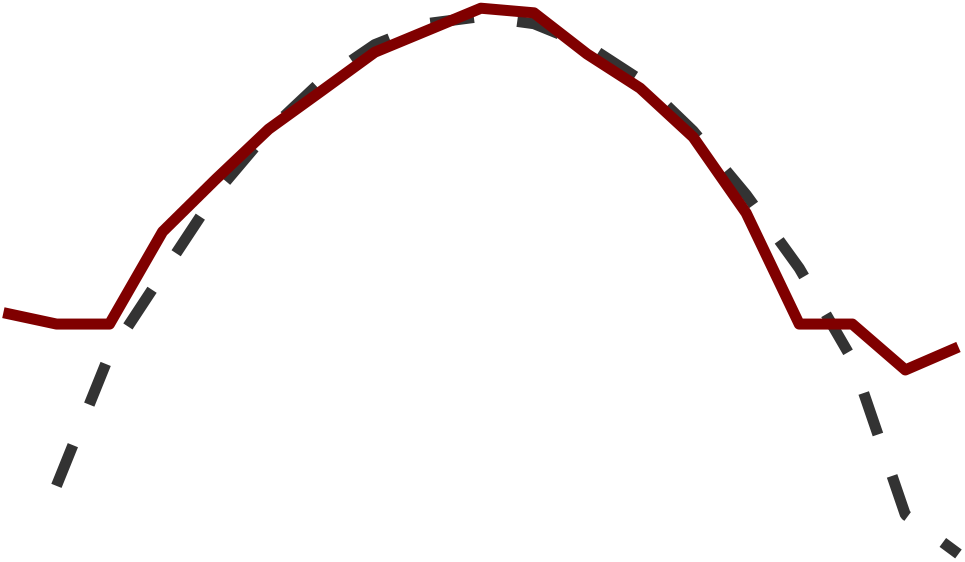
Sample patient’s forcing statistics. The red curve shows the distribution of the forcing (*V*_11_) in manuscript Figure 3. The black curve shows the Gaussian distribution of random variables with the same standard deviation as *V*_11_. The distribution of the forcing exhibits a nearly symmetrical pattern with heavy tails, indicating the presence of infrequent and extreme forcing events.

**Figure 5:**
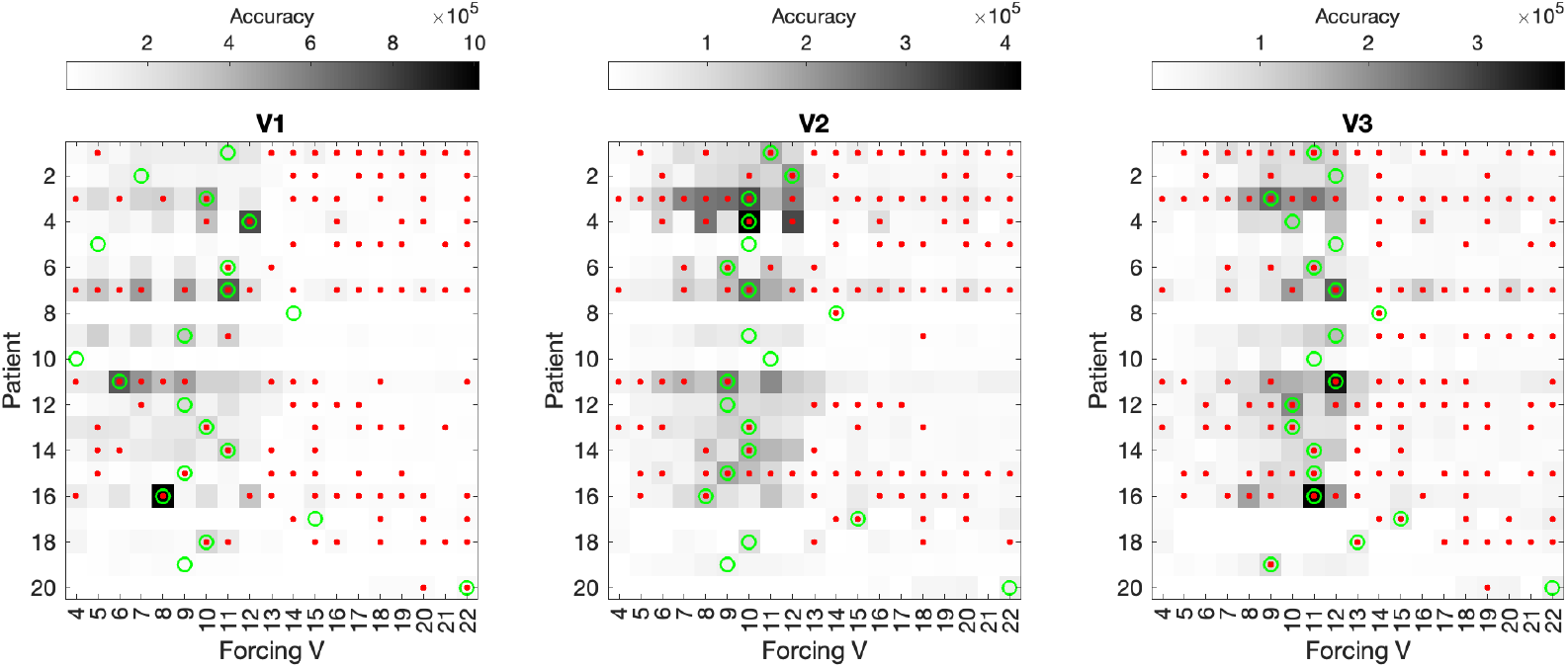
Accuracy of the Linear system of IEA delay-embedded coordinates’ predicted trajectory *d*. The accuracy (the inverse of mean squared error) of the predicted first three leading coordinate (*V*_1_, *V*_2_, and *V*_3_) time series for systems of different sizes with *V*_4_ to *V*_22_ as forcing. Red dots indicate significantly higher accuracy than those of the future trajectory predictions of the phase-randomized IEA-count null time series (*p* < 0.05, two-tailed, *n* = 50 iterations, FDR corrected for multiple comparisons across patients and delay-embedded coordinates).

**Figure 6:**
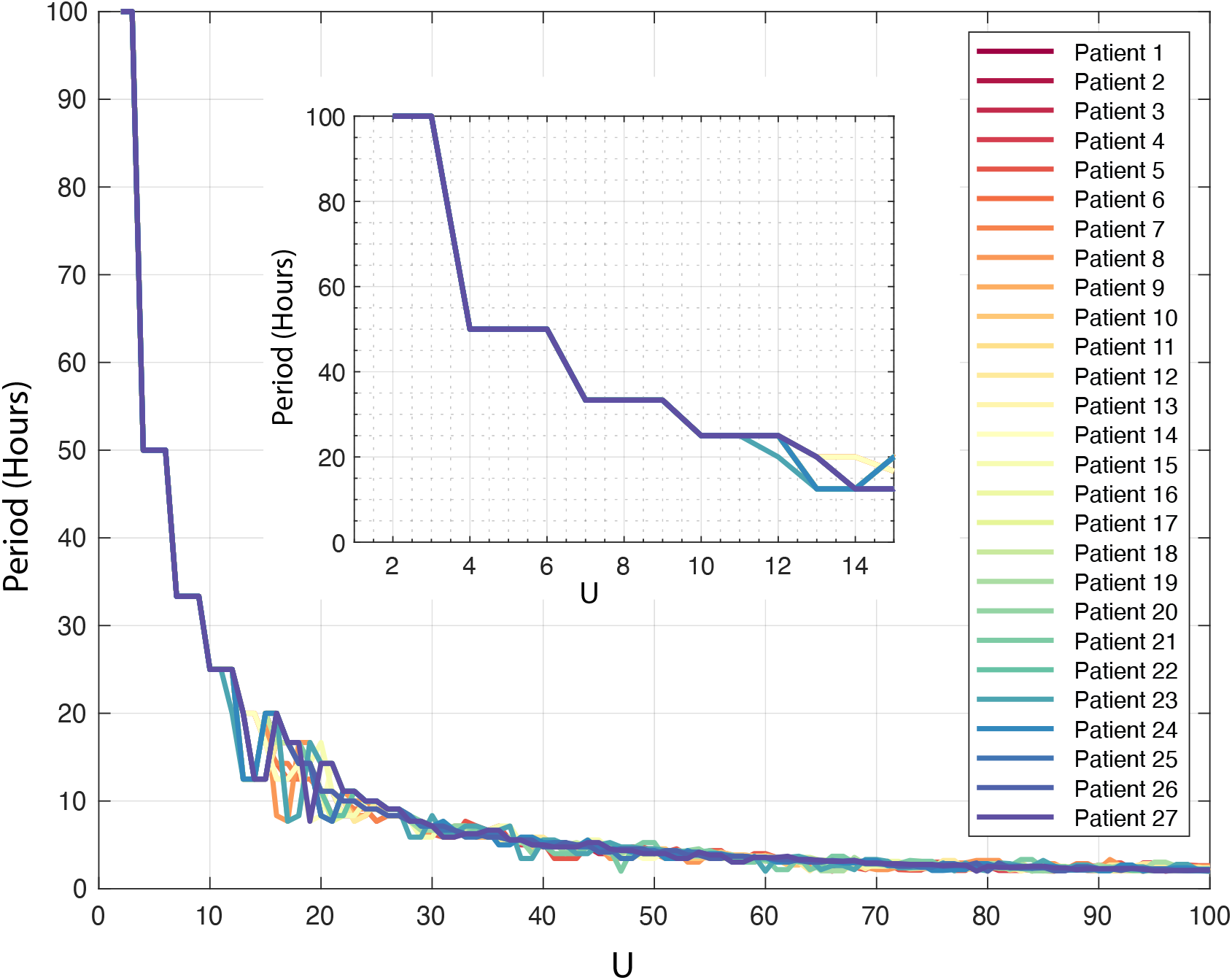
Peak periods of the *U* basis vectors. The period of peak frequency identified for all *U* basis vectors identified using Fast Fourier Transform (FFT). Different Patients are color-coded. Inset shows the same plot for *U*_1_ and *U*_15_ values.

**Figure 7:**
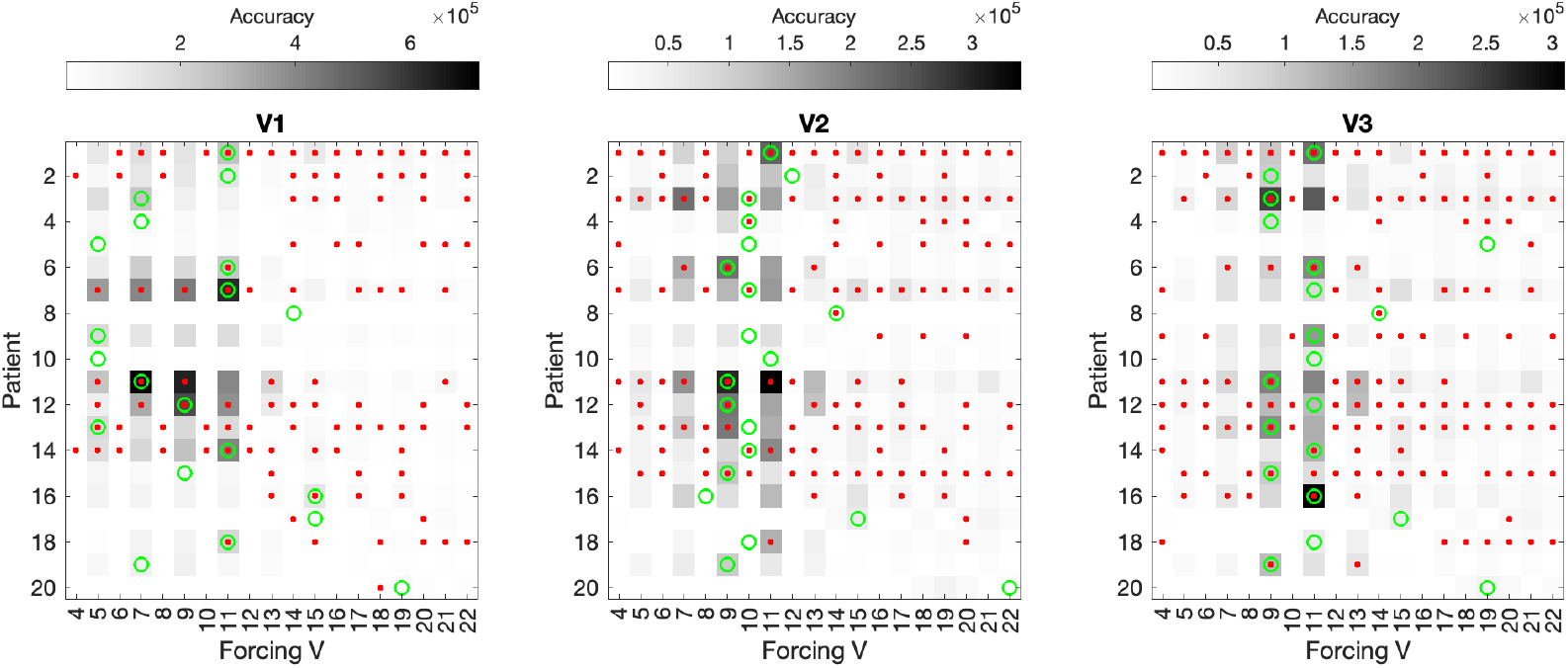
Accuracy of the Linear system of IEA delay-embedded coordinates’ predicted trajectory using convolution-based forcing calculation *d*. The accuracy (the inverse of mean squared error) of the predicted first three leading coordinate (*V*_1_, *V*_2_, and *V*_3_) time series for systems of different sizes with *V*_4_ to *V*_8_ as forcing. The forcing was directly calculated from the IEA-count time series by convolution with forcing’s corresponding basis vector (i.e., *U*_*r*_ column of *U* matrix). Interestingly, the accuracy of the systems with an odd number of dimensions is lower due to the inverted sign of the predicted time series (i.e., the predicted time series are flipped compared to the original delay-embedded coordinates’ trajectories).Red dots indicate significantly higher accuracy than those of the future trajectory predictions of the phase-randomized IEA-count null time series (*p* < 0.05, two-tailed, *n* = 50 iterations, FDR corrected for multiple comparisons across patients and delay-embedded coordinates).

**Figure 8:**
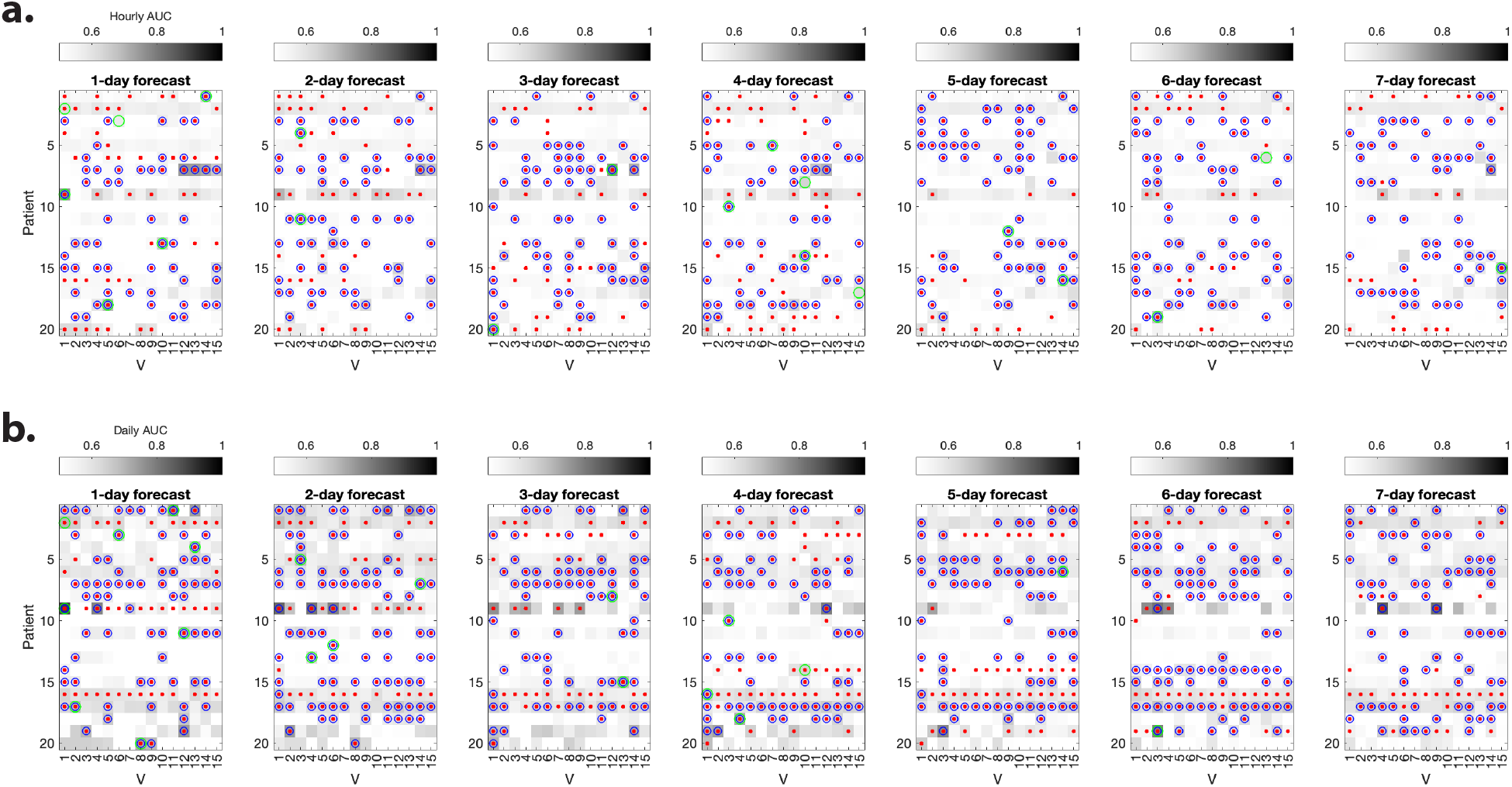
Delay-embedded coordinates of the IEA-count time series enable forecast of the seizure risk. The mean hourly ***(a)*** and daily ***(b)*** Area Under the Receiver Operating Characteristic Curve (AUC) for the forecast (1-to 7-days) of seizure risk (long IEA events) using different delay-embedding coordinates over 50 repetitions of the analysis (see Materials and Methods for classification details). Red dots show the mean AUC values that are significantly higher than those of the random null forecast (two-sample *t*−test, *p* < 0.05, Bonferroni corrected for multiple comparisons across patients and coordinates. See Statistics section for more details on the null and permutation test). Blue circles show the mean AUC values that are significantly (two-sample *t*−test, *p* < 0.05, Bonferroni corrected for multiple comparisons) higher than the AUC values calculated from the two slow peak features (i.e., their amplitude and phase). Green circles show the maximum mean AUC across all coordinates and all days for each patient.

**Figure 9:**
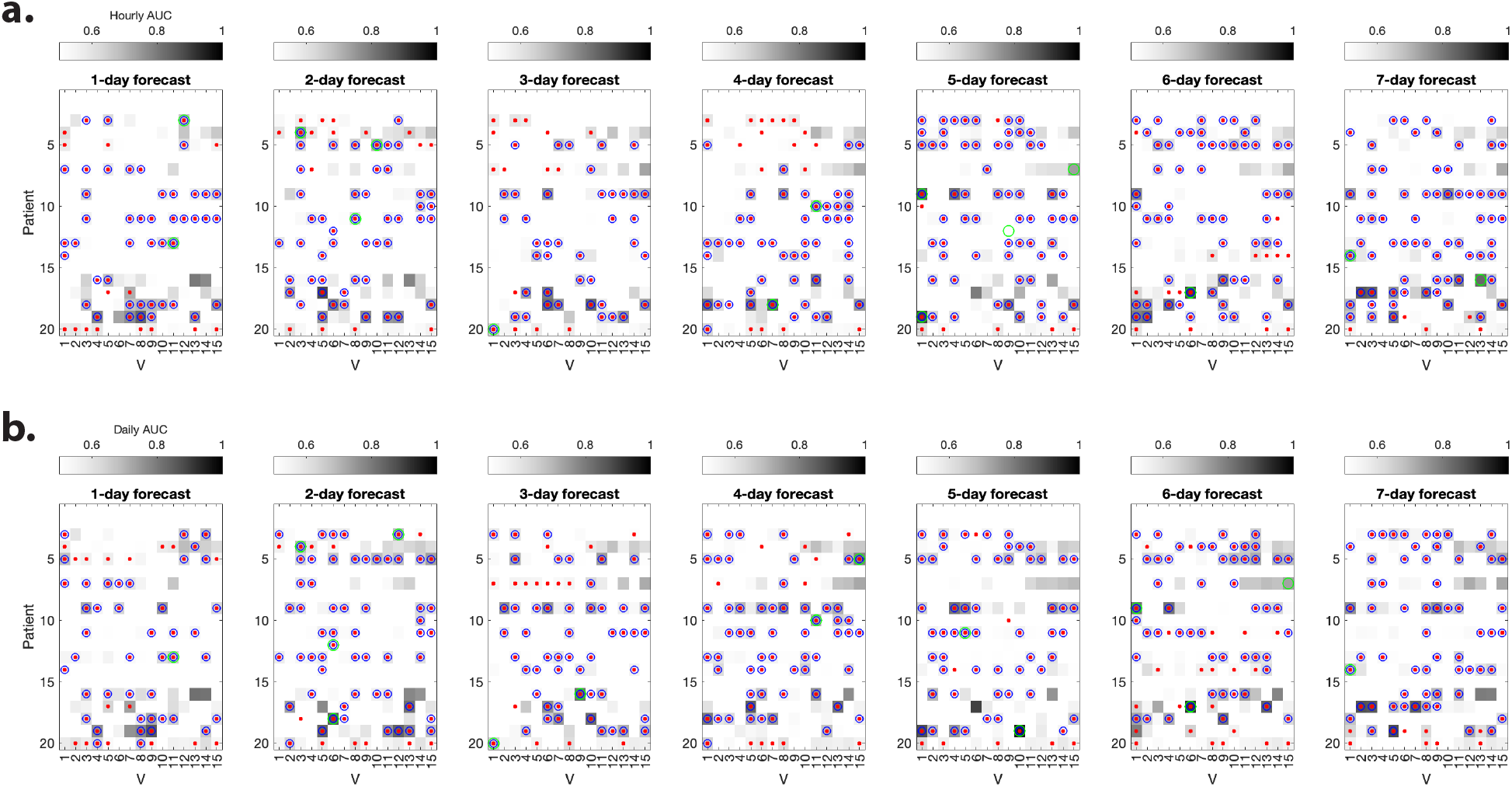
Delay-embedded coordinates of the IEA-count time series enable forecast of the patient-labeled seizure risk. The mean hourly ***(a)*** and daily ***(b)*** Area Under the Receiver Operating Characteristic Curve (AUC) for the forecast (1-to 7-days) of patient-labeled seizure risk using different delay-embedding coordinates over 50 repetitions of the analysis (see Materials and Methods for classification details). Red dots show the mean AUC values that are significantly higher than those of the random null forecast (two-sample *t*−test, *p* < 0.05, Bonferroni corrected for multiple comparisons across patients and coordinates. See Statistics section for more details on the null and permutation test). Blue circles show the mean AUC values that are significantly (two-sample *t*−test, *p* < 0.05, Bonferroni corrected for multiple comparisons) higher than the AUC values calculated from the two slow peak features (i.e., their amplitude and phase). Green circles show the maximum mean AUC across all coordinates and all days for each patient.

**Figure 10:**
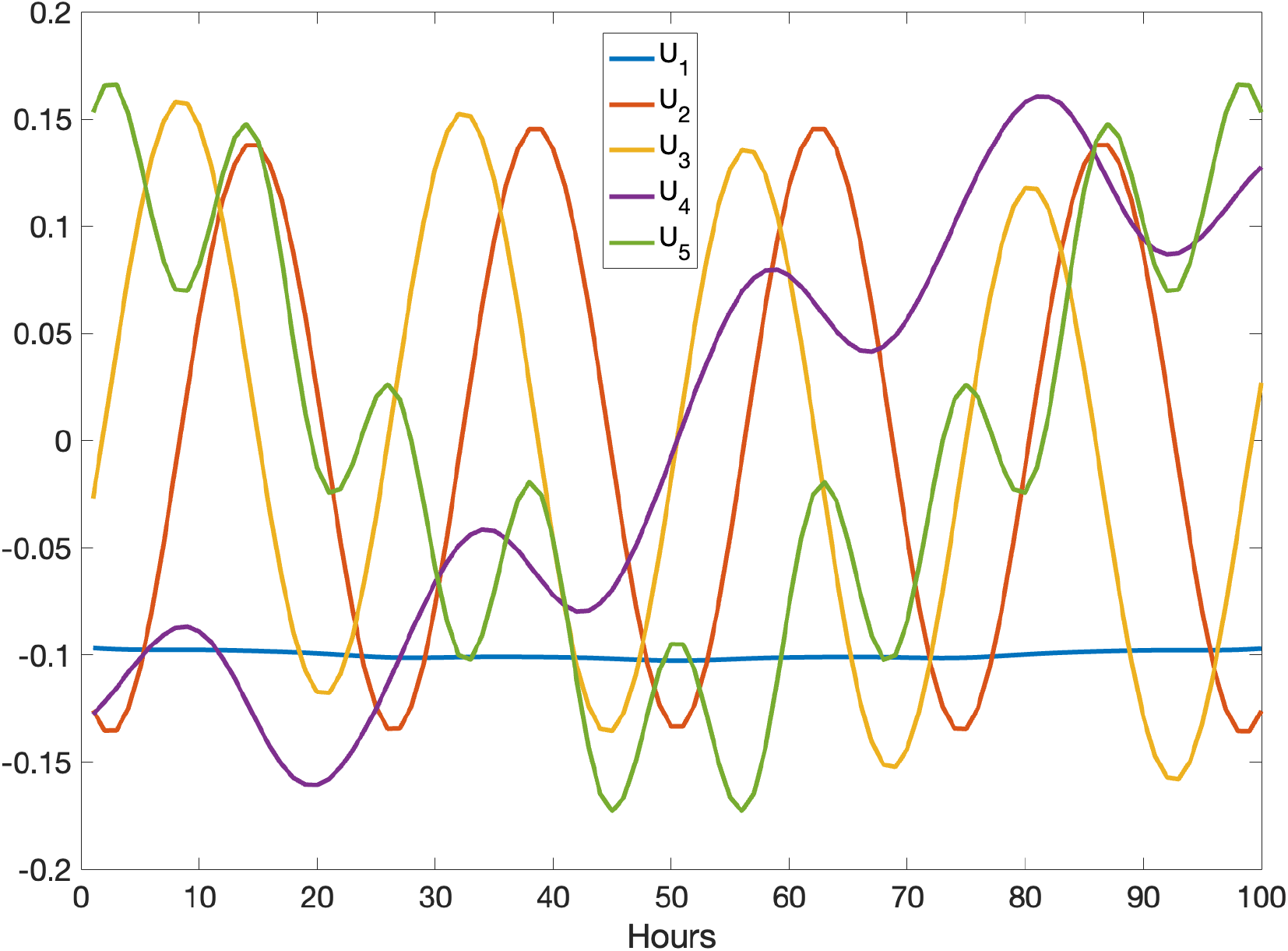
The *U* basis vectors 1 to 5 without low-pass (3-day) IEA-count time series filtering.

## Notes

### Competing Interest Statement

The authors have declared no competing interest.

